# Comparative pangenome analysis of major pneumococcal genotypes from India

**DOI:** 10.1101/2024.01.14.575557

**Authors:** Sreeram Chandra Murthy Peela, Sujatha Sistla

## Abstract

**Background:** Pneumococcal genomes are highly dynamic with varying core genome sizes. The genotype classification system, Global Pneumococcal Sequence Clusters, identified patterns within genotype and antibiotic resistance. Few genotypes like GPSC10 are frequently associated with antimicrobial resistance and high rates of non-vaccine serotypes.

**Objective:** To identify and annotate the differences in the core genomes of major GPSC in India, and construct and analyse the Indian Pneumococcal Pangenome (IPPG).

**Methods:** Using existing dataset from the Global Pneumococcal Sequencing Project, 618 strains were included. The most frequent GPSCs: GPSC1, GOSC2, GPSC8, GPSC9 and GPSC10 were analyzed separately. Pangenomes were constructed using Panaroo with tuning the family threshold parameter. Differences in protein clusters were identified using Orthovenn3 webserver. Functional annotations were performed by eggNOG, Uniprot and STRING database searches.

**Results:** The IPPG core genome size (1615 genes) was similar to those reported previously, with similar distribution of metabolic categories across the five GPSC types. The GPSC10 (1619 genes) and GPSC1 (1909 genes) had the lowest and highest core genome sizes respectively, and these core genomes possessed genes encoding for macrolide and tetracycline resistance. Virulence genes ply, psaA, pce (cbpE), pavA, nanB, lytA, and hysA are detected among all the core genomes.

**Conclusions:** There is a genotype specific variation within the core genomes of major GPSCs in India. The presence of antibiotic resistance genes among GPSC1 and GPSC10 core genomes explain widespread drug resistance due to these genotypes. The core virulence genes identified among all the genotypes indicate conserved pathogenesis mechanisms, and can be targets for vaccine development or therapy.

## Introduction

*Streptococcus pneumoniae* is a Gram-positive pathogen responsible for a variety of infections in young children and old adults. Owing to the high burden of pneumococcal infections, the Government of India introduced pneumococcal conjugate vaccine in the National Immunisation programme in 2021 in a phased manner [1]. Ongoing surveillance of pneumococcal serotypes is primarily performed by the Global Pneumococcal Sequence (GPS) project, wherein the serotypes, antimicrobial resistance mechanisms and genomic diversity are studied [2]. Currently, the second phase of the project began, and the data from the first phase (until 2018) was deposited in the GPS project database. Within this database, the metadata and annotated genomes are available for public download and reuse. From India, 618 pneumococcal isolates from both diseased and asymptomatic carriers were collected and analyzed.

Based on the global estimates and analysis of Indian data, a few genotypes, designated as the Global Pneumococcal Sequence Clusters (GPSCs), are associated with drug resistance and non-vaccine serotypes [3]. The GPSC10 is predominantly associated with non-vaccine serotypes, and contains the CC230 clonal complex that emerged as the multidrug resistant clone post-PCV10 introduction in USA. Recently, the GPSC10-24F is identified as the most prevalent non-vaccine type with high rates of multidrug resistance [4]. The GPSCs are also associated with various metabolic genotypes (MT-types), indicating a difference in metabolic control which can impact the strain replacement when under competition from a different MT type [5].

The genome of pneumococcus is highly dynamic with high rates of natural transformation. Traditional genome analysis focused on mapping reads to a reference genome (usually the TIGR4 strain), and therefore the full genomic variants were not analyzed. On the other hand, the use of pangenome, the total gene content of a species, is gaining importance in comparative analysis [6]. Using a pangenome as a reference, the rare genes and associated variants can be identified. Construction of pangenome typically involves graph-based algorithms, and generally use genome annotation files and genome aligners like MAFFT for constructing a core genome alignment.

To specifically understand the variations among the major GPSC types and to identify patterns within the genetic makeup of these GPSC types, an exploratory study was designed to study pangenomes of pneumococci isolated from India. Using a combination of pangenome functional annotation and comparative proteome analysis, we aimed to identify the differences within the core genomes of the major GPSC types and the Indian Pneumococcal Pan-Genome (IPPG) of the 618 isolates.

## Methods

### Genome data

The publicly available annotated pneumococcal genomes (N=618) collected through the GPS project in India were included in the study. The genome sequences, annotations and metadata were downloaded from GPS project database available at https://data.monocle.sanger.ac.uk/ (last accessed on 01 June 2023). Most of the isolates were collected during pre-PCV period (prior to 2017) while 34 isolates were collected in 2018 (the post-PCV period).

### Pangenome construction

The pangenomes were constructed using Panaroo v1.2.10 (available as a docker image) [6]. From the 618 genomes, 100 genomes were randomly selected and pangenome was constructed using a range of family thresholds (0.7, 0.8, 0.9, 0.95, 0.98, and 1). Using these thresholds, and setting all other parameters to default values, pangenomes were constructed from these test data. The genome compartments were designated by setting 95% as the threshold for number of isolates to group in the core genome compartment. Using scree plot, the highest family threshold with highest core genome size (number of genes identified in core genome) was selected for subsequent analysis (Supplementary Figure 1). From initial visualization, the optimum family threshold was identified as 0.95.

**Supplementary figure 1:**
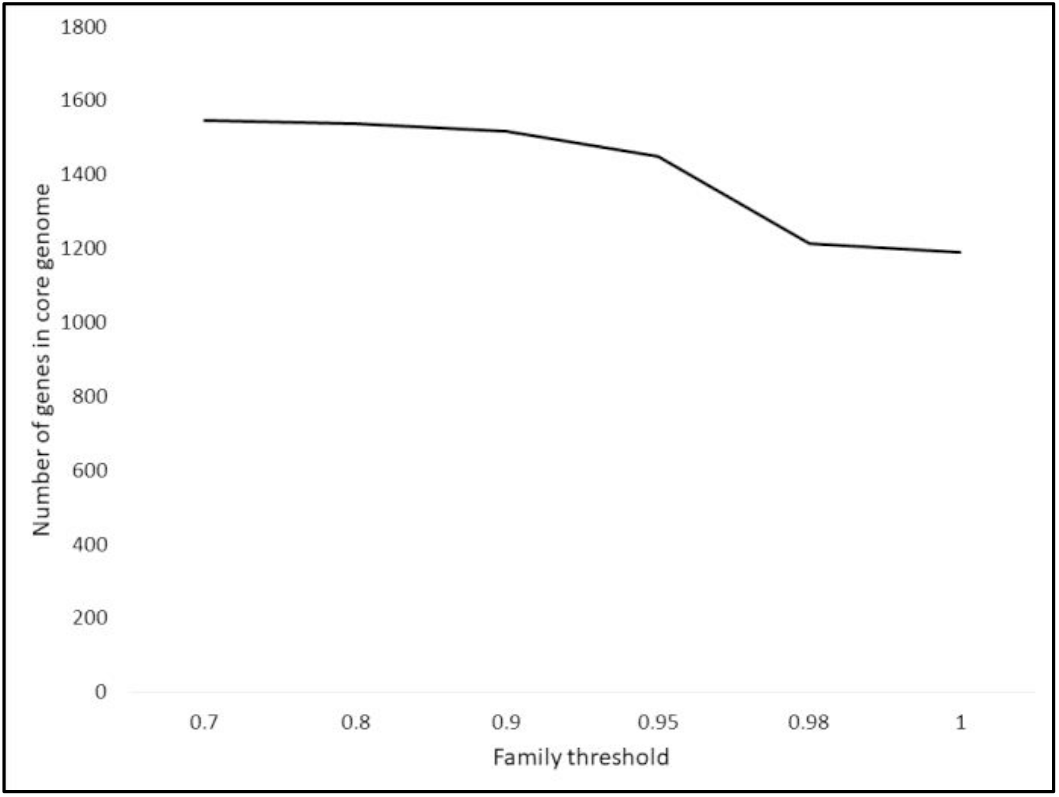
Change in core genome size with different family thresholds. At a threshold of >0.95 core genome size reduced significantly.

Using this family threshold, pangenome was constructed for all the isolates (hereafter referred as”Indian Pneumococcal Pan Genome, IPPG”). The most prevalent five GPSCs (GPSC1, GPSC2, GPSC8, GPSC9, and GPSC10) were analyzed separately by constructing pangenomes for each of these GPSCs. For the construction, strict mode with removal of invalid genes, and core genome alignment with mafft were selected. The pangenome construction was carried out in a workstation with 10 cores of i7-11800H (2.30 GHz) and 32 GB RAM, and approximately 24 hours were taken for constructing IPPG, while for each GPSC, the construction completed in less than two hours depending on the number of samples.

### Phylogenetic analysis

Previous study using similar data employed reference genome mapping and creating phylogenetic tree using the identified variants. In this study, both core genome alignment and gene presence-absence file from Panaroo were used. We specifically employed this method to use variant information that may be absent when using a single reference genome. The phylogenetic trees were constructed separately with IQ-TREE v1.6.1 using core genome alignment file (variants extracted using SNP-SITES v2.3.3) and gene presence-absence file (pseudo-FASTA file was generated by categorical encoding of presence or absence of a gene in a sample using customized Python script) [7,8]. Bootstraps of 1000 and auto model selection were used for both tree constructions, and the resulting phylogenetic trees were merged later using IQ-TREE consensus tree construction. The tree was annotated and visualized in iToL v6.

### Core genome annotation

From each GPSC pangenome and IPPG, using the gene presence-absence file, core genes were extracted using a customized Python script. The resulting multi-FASTA files were then re-annotated using Prokka v1.14.6 [9]. The non-coding RNA annotation was facilitated by using”--rfam” option while running Prokka. From the annotated core proteomes, orthologs were identified using Orthovenn3 web application [10]. The annotated core proteomes of major GPSCs and IPPG were searched using eggNOG database version 5 using the web-based e-mapper tool and setting the following parameters: E-value = 0.00001, minimum identity, query and subject coverages set to 80%, enabling SMART annotation, and taxonomic scope = 2 (bacteria) [11]. The antibiotic resistance and virulence genes were identified using Abricate v1.0.1 and VFDB and NCBI AMR databases (versions accessed on 16-11-2022) [12–14]. The virulence genes associated with streptococci were downloaded from VFDB, and GO annotations were retrieved using eggNOG and emapper as described above. From this search, 222 unique GO terms were identified for the virulence genes. Using the GO annotations of these virulence factors and one-to-one mapping, the probable virulence genes and their GO terms were identified for GPSC types.

The relative enrichment of identified gene ontology terms with respect to the whole genome (R6 strain) was analyzed using STRING functional enrichment application stringApp v2.0.2 in Cytoscape v3.10.1 [15]. For importing the network into Cytoscape, the STRING ids retrieved from the eggNOG database search was used, and only those proteins identified to be from R6 (taxon id: 171101) were included. For each GO annotation term, the average number of genes across all GPSC (and IPPG) types was calculated by including the number of background genes, and Z-scores for each GPSC (and IPPG) type were calculated using customized R script. All visualizations were performed either in MS EXCEL or R v4.1.3 (using ggplot2 library).

## Data availability

The genome annotations and metadata are available in the GPS project database (https://data.monocle.sanger.ac.uk/). The samples selected for analysis including their metadata is available in the supplementary file. The core proteome files and the supplementary files are deposited in FigShare (https://figshare.com/articles/dataset/GPSC_pangenome_analysis/24993954).

## Results

### Pangenome comparisons

Together the five GPSCs accounted for more than one-third (205/618) of the isolates. The Indian pneumococcal pangenome had a total of 4919 genes with 1534 and 81 genes within the core (detected in >99% of the isolates) and soft-core (detected in 95-99% of the isolates) compartments. The core genome size was lowest for GPSC10 (1694 genes) and highest in GPSC1 (1909 genes) (Figure 1). Although the number of isolates for GPSC8 was the least, its core genome size was similar to GPSC1 (Table 1).

**Table 1:** Core genome sizes and number of samples analyzed for each GPSC type.

| GPSC type | No. of isolates | Core genome size (core + soft core) |
| --- | --- | --- |
| GPSC10 | 83 | 1694 |
| GPSC9 | 29 | 1790 |
| GPSC2 | 21 | 1788 |
| GPSC1 | 52 | 1909 |
| GPSC8 | 20 | 1907 |

**Figure 1:**
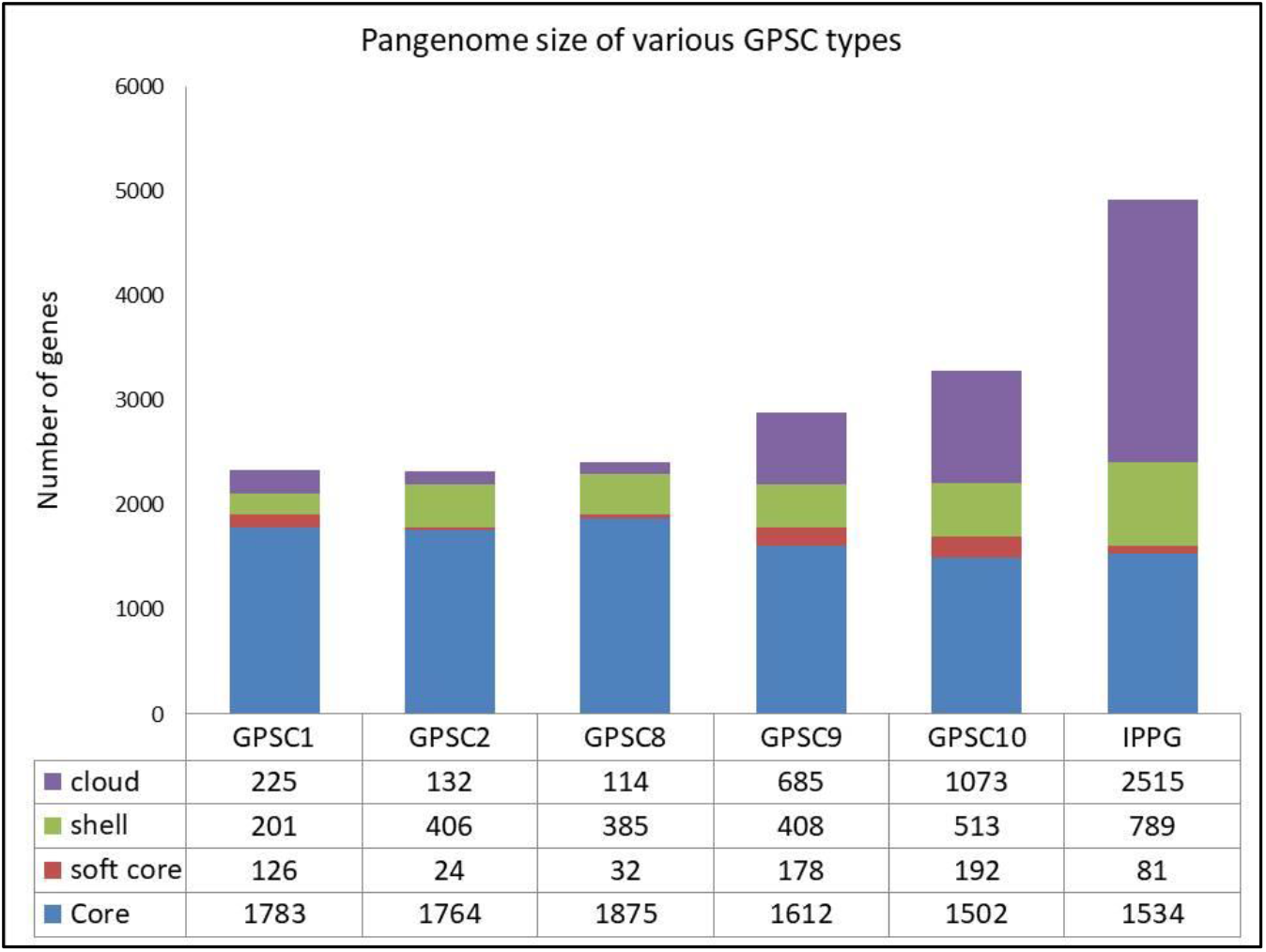
Core genome and other pangenome compartment sizes among the five GPSC (and IPPG) types. The Indian pneumococcal pangenome (IPPG) had the highest number of genes in its cloud compartment.

Interestingly, a negative trend was observed – as the number of samples increase, the core genome size reduced (Supplementary Figure 2).

**Supplementary figure 2:**
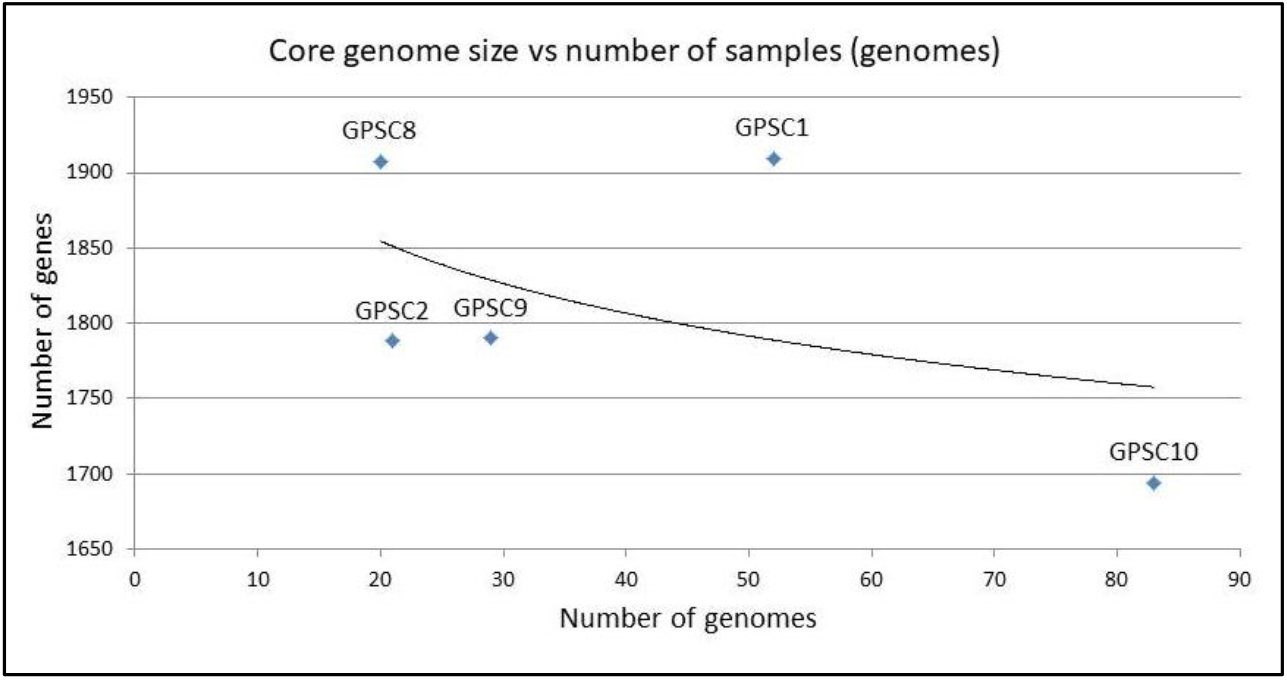
Trend in core genome size compared with the number of genomes. The trend line was calculated using MS EXCEL power regression type, and has R^2^ = 0.2112

### Phylogenetic relationship

Using a combination of input parameters and generating a consensus tree, the isolates GPSC types were identified within similar clades (Figure 2). Most of the non-vaccine serotypes (blue color in the figure) were also clonally related. It is interesting to note that the GPSC10 isolates were grouped into two different clades, and most of the non-vaccine serotypes were associated with a single clade.

**Figure 2:**
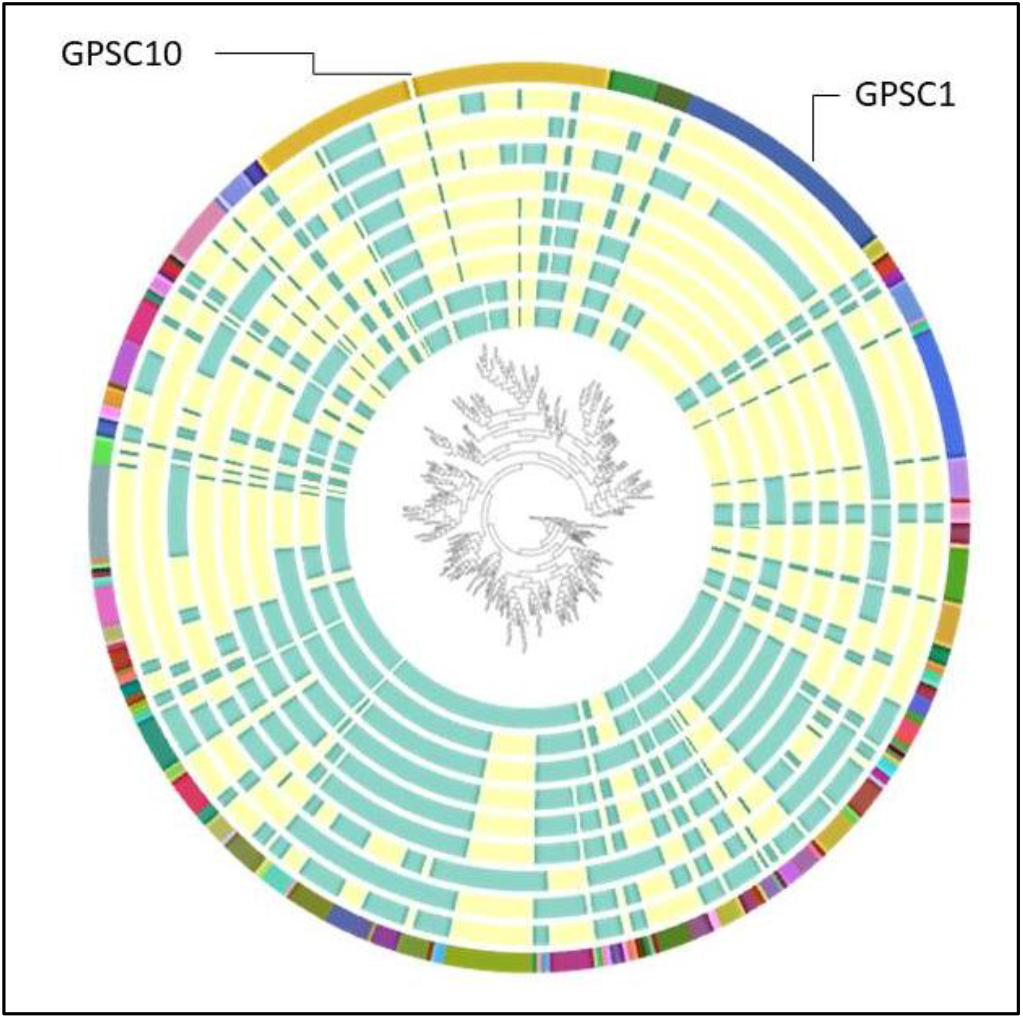
Phylogenetic relationship among the 618 isolates. The strips (from inside) represent vaccine coverage (yellow in color) for vaccines PCV7, PCV10 (GSK), PCV10 (Pneumosil), PCV13, PCV15, PCV20, PCV21, PCV24, and IVT-25. For visualization, the two most common GPSC types were labeled. The full file with metadata can be viewed at https://microreact.org/project/muCTm4owiDE51AaJuNSM6H-indiacoregenome619

### Core genome annotations

Among the core proteomes, the least number of proteins identified in the category S (Unknown function) were detected in GPSC10 (93 genes when compared with an average of 247.3 genes for all GPSCs) (Figure 3). Only one protein under category Z (cytoskeleton) was identified only within GPSC2. None of the core genomes had COG categories for B (chromatin structure and dynamics), W (extracellular structures), X (mobilome: prophages, transposons) and Y (nuclear structure). The GPSC10 had slightly higher number of genes associated with category Q (Secondary metabolite biosynthesis, transport and catabolism) while in GPSC8, category V (defense mechanisms) had slightly higher number of genes when compared with average number of genes across each category.

**Figure 3:**
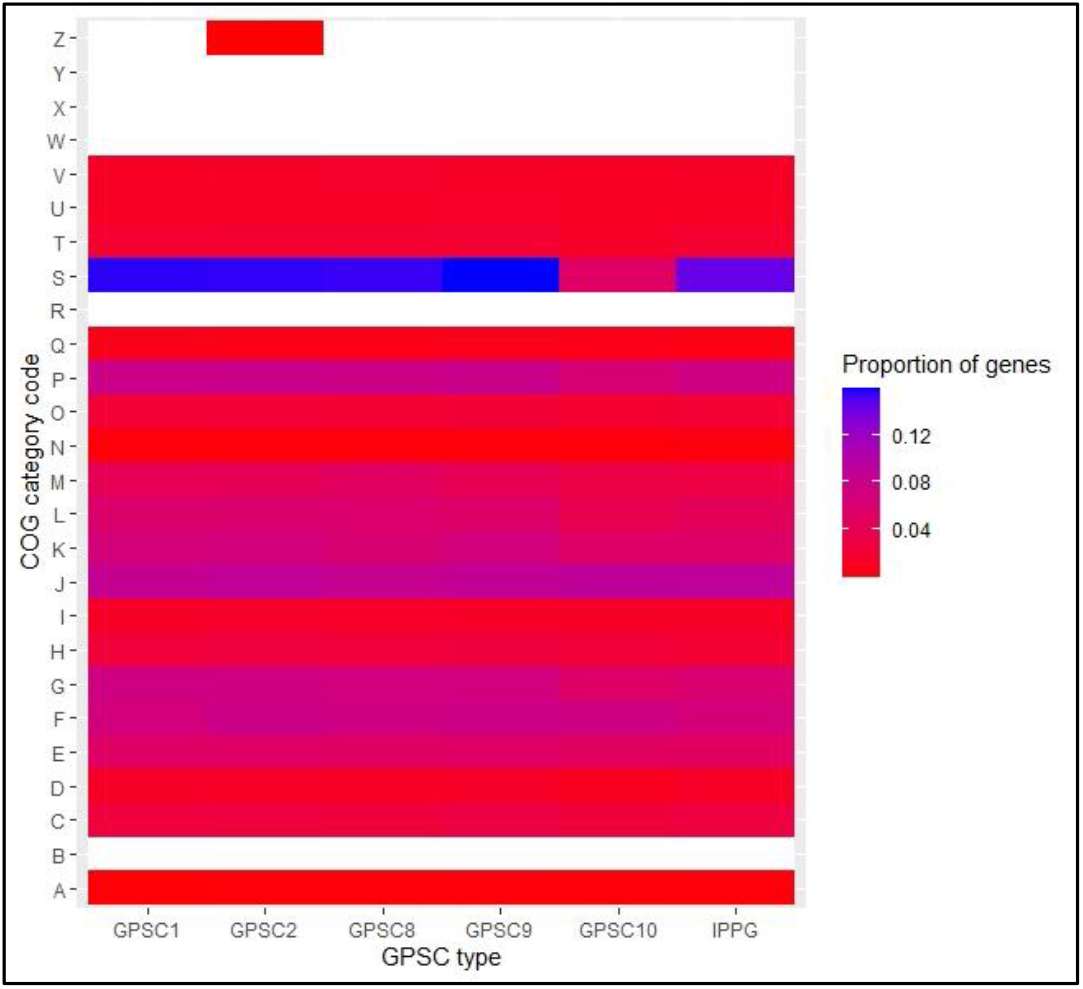
Various COG categories among the different core genomes. The categories were obtained by using eggNOG-emapper v2.1.12. The colors represent proportion of genes in each category. Raw counts are available in the supplementary file.

Among the IPPG core genome, no antibiotic resistance genes and 10 virulence genes (PavA, Ply, PsaA, LytC, Pce, HysA, CbpD, PfbA, NanB, and PavB) were detected, and had the least number of virulence genes (Figure 4). The GPSC1 core genome had the highest number of virulence genes, and a few genes (cbpA in GPSC1, for example) had allelic variants. The IPPG, GPSC2, GPSC8, GPSC9 core genomes had no antimicrobial resistance genes, while tetM was detected in both GPSC10 and GPSC1, and mefA-msrD in GPSC1 core genome alone. Among the 10 identified virulence genes within IPPG-core genome, five were annotated by the emapper tool. These belonged to various categories (M – 2, one each of S, G and P), indicating cell wall/membrane/envelope biosynthesis or inorganic ion or carbohydrate transport or metabolism. Among the virulence genes unique for GPSC1, only sortases (srtBCD) were annotated by the emapper.

**Figure 4:**
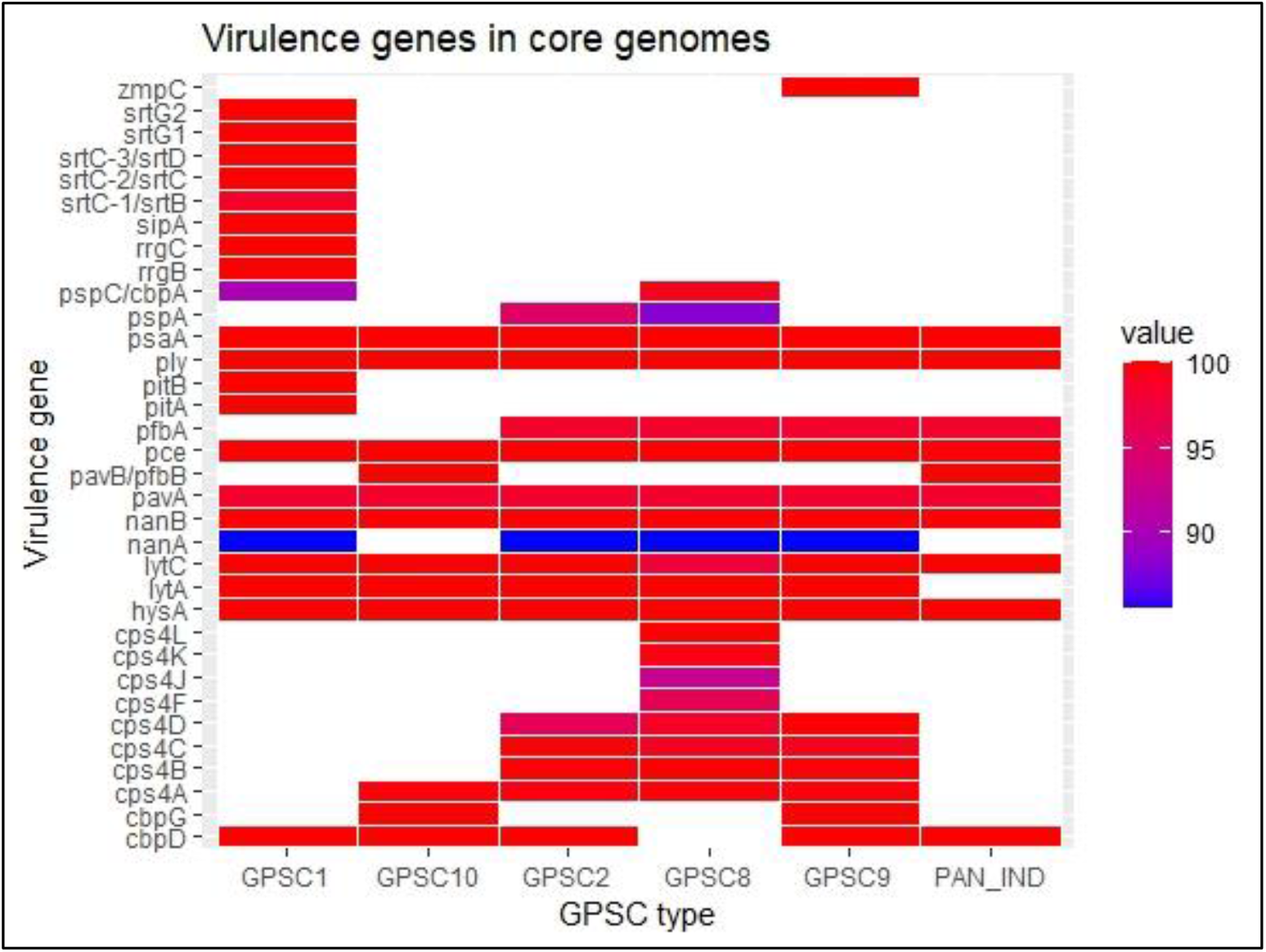
Distribution of various virulence genes in the core genomes of five major GPSCs and the Indian Pneumococcal Pangenome (PAN_IND). The percent identity is encoded in various colors, while white color indicates the absence of a gene.

Using STRING database search, 50 GO terms were detected among all the GPSC types and IPPG, and 88 GO terms were identified exclusive to GPSC10 core genome (full data in Supplementary File). The IPPG core genome had additional two GO terms GO: 0043043 (Peptide biosynthetic process) and GO: 0051179 (Localization). Using the average proportion of genes present among the core genomes, the relative difference in genes associated with each GO term was analyzed (Figure 5). Comparatively, higher proportion of genes were identified in GPSC8 core genome (78.6%-96.9%), while least number of genes were detected among the IPPG core genome (63%-89.5%) (Supplementary data).

**Figure 5:**
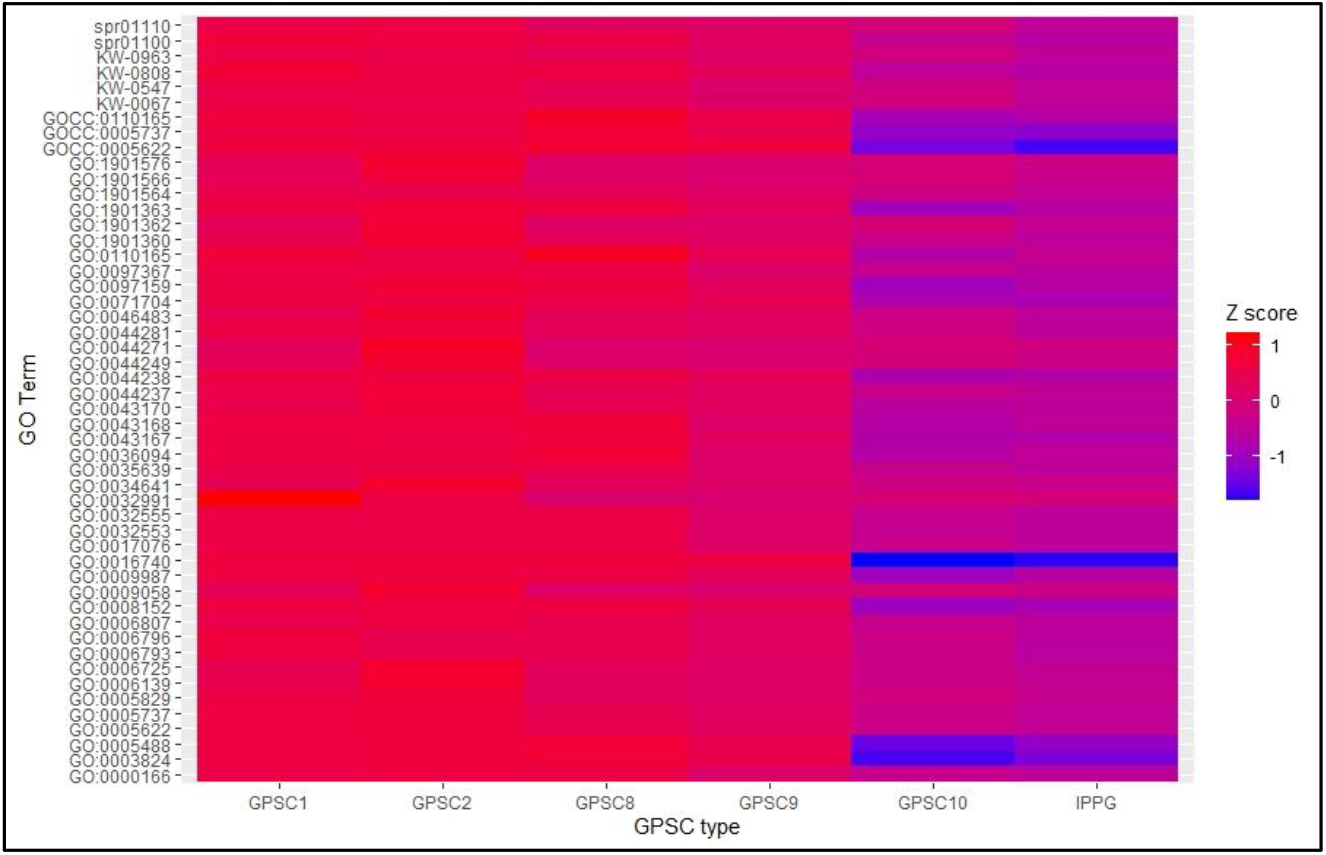
Distribution of genes detected for each GO term among the core genomes. The data is scaled by calculating the Z score of each GO term (average = number of genes identified/number of genes in background genome).

Among the GO terms unique to GPSC10, all the genes associated with ribosomal subunit (STRING cluster CL: 96) were detected, and for the remaining GO terms, the GPSC10 carried 65.7-97.8% of the background genes. Ten GO terms were associated with virulence genes, and GO: 0016787 (Hydrolase activity), GO:0071840 (Cellular component organization) and GO:0016043 (Cellular component organization or biogenesis) are associated with highest number of virulence genes (Figure 6). All these three GOs contain lytA and cpsB within the set of genes, and speB, vpr, nanB, and zmpC are the additional genes associated with GO: 0016787.

**Figure 6:**
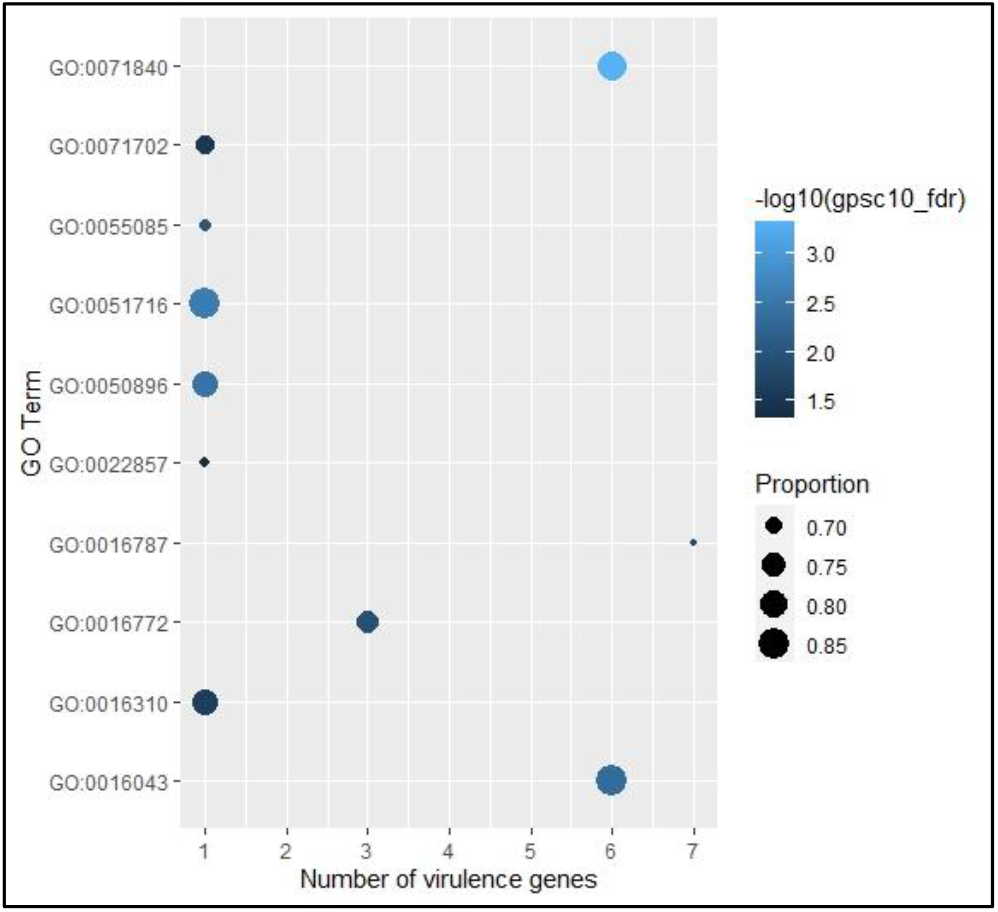
The GO terms uniquely associated with GPSC10 and number of virulence genes within each GO term. The bubbles are colored according to log10 scaled FDR values, and sized according to the proportion of genes when compared to background (total genome).

### Core genome protein clusters

Clustering the proteomes using OrthoMCL in Orthovenn3 webserver identified 902 (53.1%) shared protein clusters among the total 1698 clusters identified, and very few clusters present exclusively in the GPSC1 and GPSC8 core genomes (Figure 7). Based on the set of protein clusters shared, a phylogenetic tree revealed closely related GPSC10 and IPPG core genome (Figure 8). Among the shared protein clusters, the GO terms metabolic processes (GO: 0008152), hydrolase (GO: 0016787) and transferase (GO: 0016740) activities, and cytoplasmic compartments (GO: 0044464) were the enriched terms (Figure 9). The GPSC9 shared highest number of clusters with GPSC1 (15 clusters) and GPSC2 (15 clusters; no cellular compartment annotations). None of these clusters had any specific GO terms at a higher proportion. However, the GPSC9 and GPSC1 shared clusters were identified with GO: 0000746 (conjugation), GO: 0009234 (menaquinone biosynthetic process), and GO: 0006313 (DNA-mediated transposition) with FDR-adjusted p-values < 0.05.

**Figure 7:**
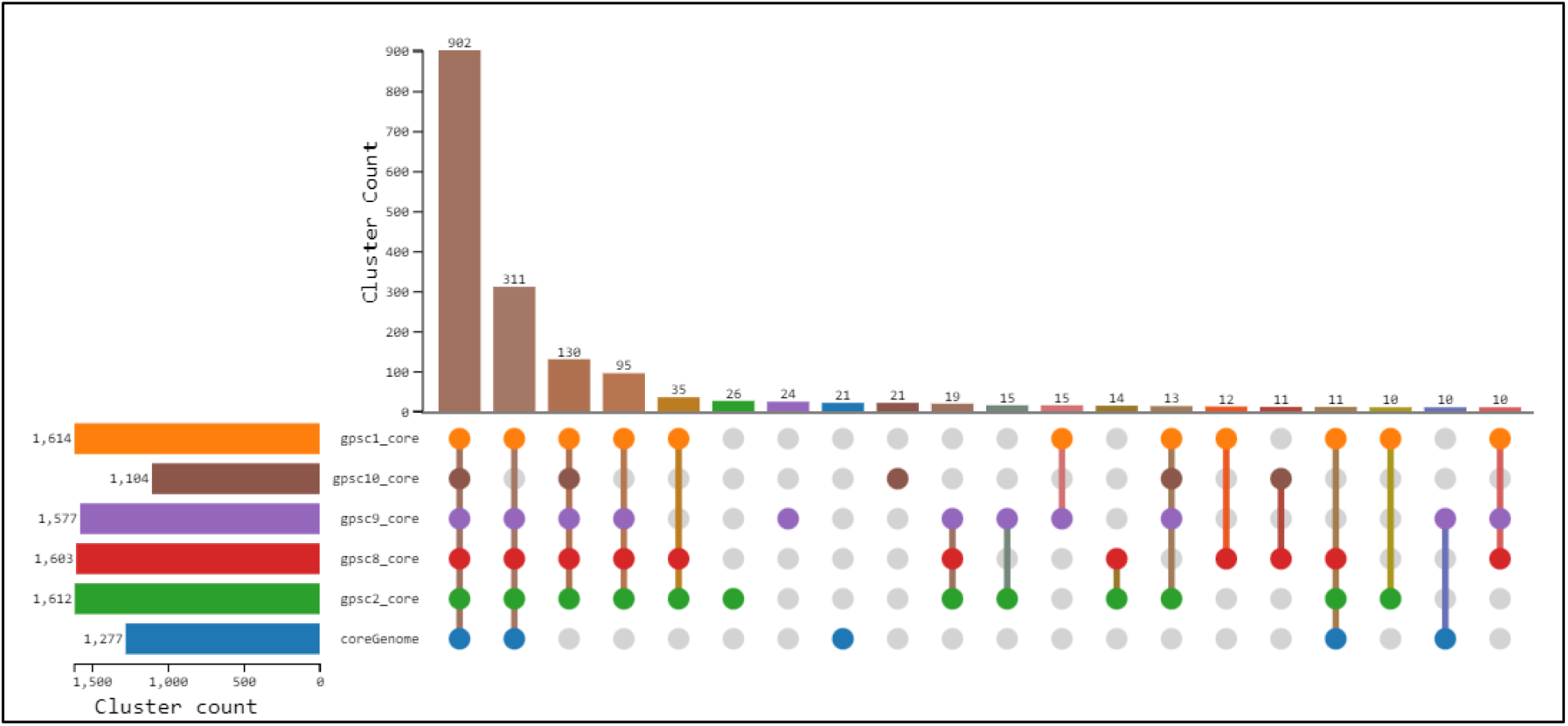
Number of shared and unique protein clusters among the core proteomes and number of proteins within each proteome. Each vertical line represents the fraction of clusters shared among the core proteomes, while unconnected dots represent unique clusters within a proteome. The Indian Pangenome is represented as”coreGenome”.

**Figure 8:**
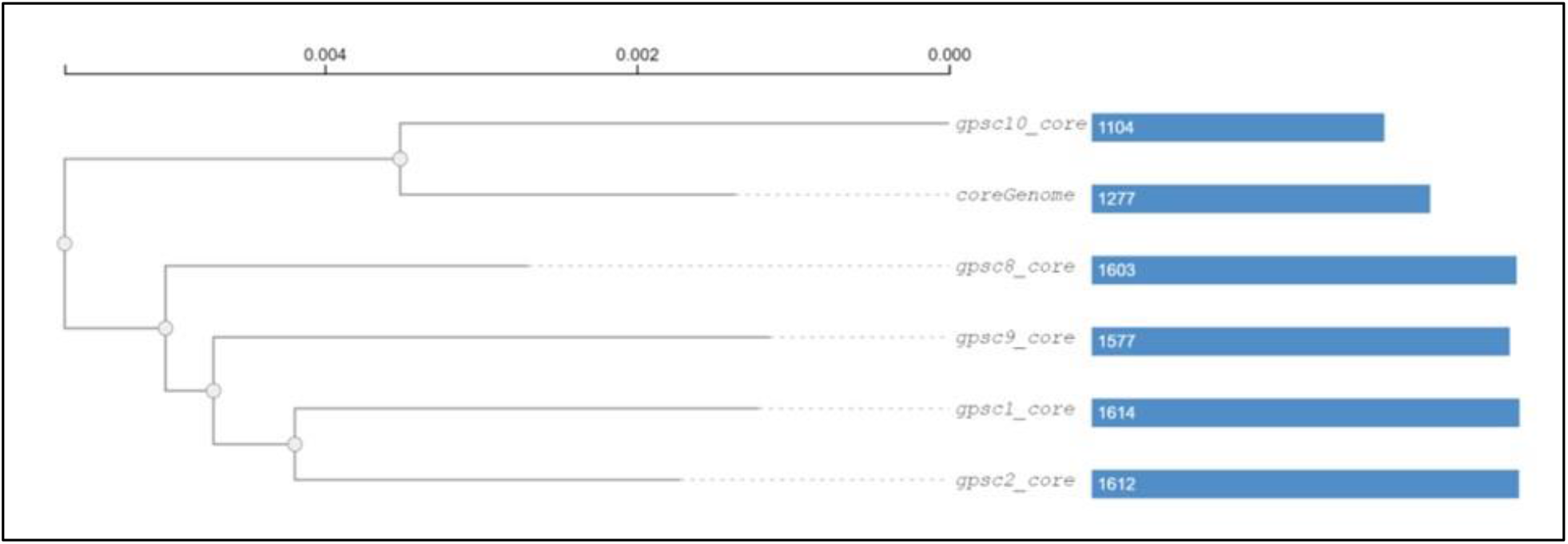
Phylogenetic tree based on the shared proteome clusters. The tree was inferred using Maximum Likelihood method with JTT+CAT substitution model in Orthovenn3.

**Figure 9:**
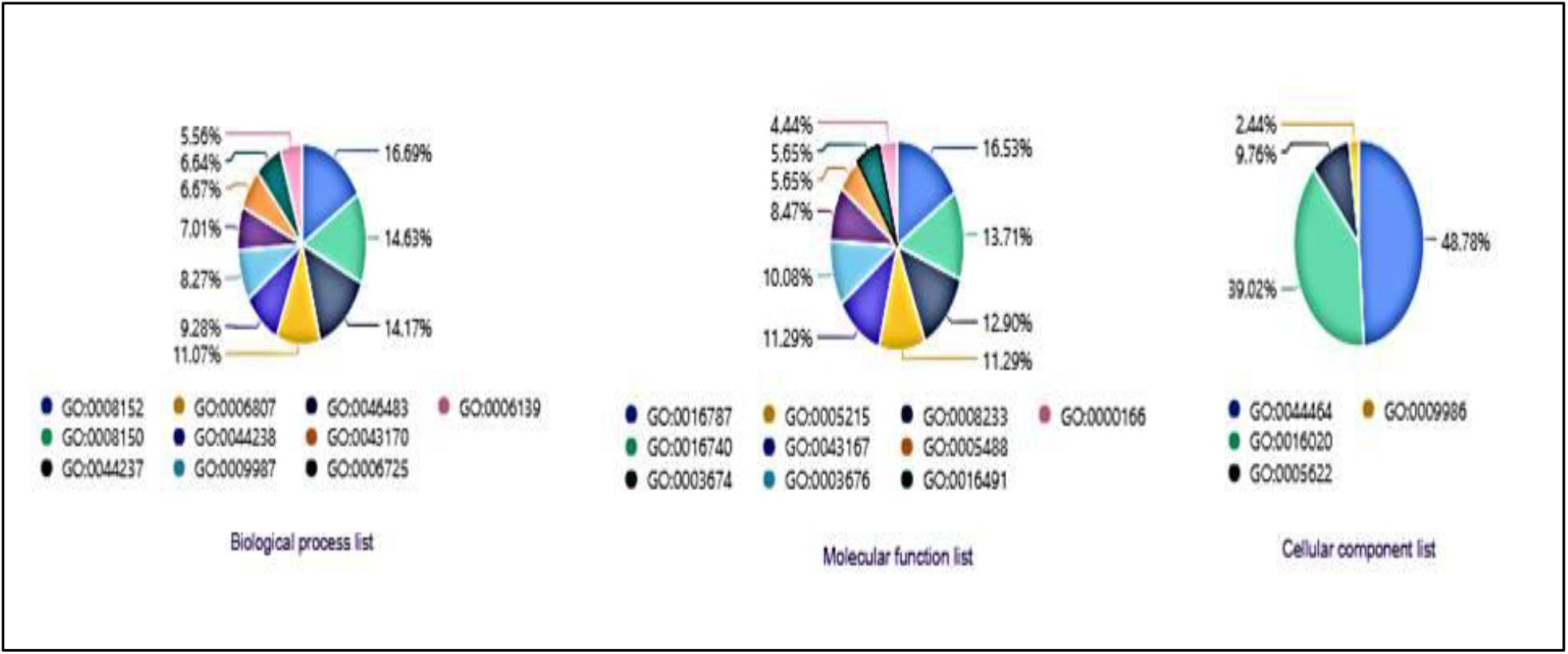
GO annotations of the 902 common protein clusters. Only the top results are represented.

## Discussion

Pneumococcal genomes are highly dynamic with continuous loss/gain of new genes, especially related to antibiotic resistance [16]. While pneumococcal vaccines are introduced as early as 2007 (in USA), in India, until 2021, pneumococcal vaccination was carried out in phased manner in select states with high burden of these infections [1]. Introducing pneumococcal vaccine in India is expected to change the local bacterial population structure as observed from multiple studies across the world. Identifying the clonal relationships among the resistant strains and non-vaccine serotypes is imperative to understand the dynamic changes in these patterns that can be associated with vaccine introduction. Based on genomic signatures, a recent classification system, termed the Global Pneumococcal Sequencing Clusters (GPSCs) emerged from the data generated during the Global Pneumococcal Sequencing (GPS) Project [3]. In India, a brief study was carried out under the aegis of the GPS project to study genomic characteristics of 480 pneumococcal strains isolated from both healthy and diseased subjects during 2009-17 (the”pre-vaccine era”) [17]. While multiple GPSC-types were observed, indicating divergence within the strains, a few GPSC types were more frequently observed, and these genotypes were associated with specific serotypes. For example, the combination GPSC1-19F (CC320) was most frequent, while GPSC10 (CC230) consisted of both vaccine and non-vaccine serotypes [17]. The GPSC10 clone is also a high risk clone with high rates of drug resistance, and is also identified as a major challenging clone for vaccine introduction [4].

The pangenome contains all the genes identified among multiple strains within a bacterial species. Multiple tools exist for pangenome construction, and these tools employ a variety of algorithms (graph-based) to deduce the pangenome. To understand the relative distribution of genes across multiple strains, the terms core genome, shell genome and cloud genome are generally used. By default, most studies considered genes present in at least 95% of the strains as the core genome, while genes in less than 15% of the strains as”cloud” genome [6]. The pneumococcal pangenome is considered to be of”open” type, and can increase in size as more number of strains is included in the analysis [16]. Analyzing pangenomes can aid in understanding the full genetic repertoire of the pathogen, and in identifying variations across the strains. Recently the pangenomes are also being recommended instead of a single reference genome to identify strain-level genomic variants. The use of pangenome-based analysis in highly dynamic pathogen like pneumococcus can therefore provide additional insights into functional and phenotypic divergence among the strains.

Among the 618 genomes analyzed using a combination of core genome phylogeny and gene presence-absence matrix, the non-vaccine serotype isolates among GPSC10 were clonally related. One immediate observation could be that these isolates are an effect of recent serotype switch rather than replacement. As evidenced from a recent study, serotype switch from 14 to 11A may have been occurring within the Indian pool of isolates [17]. However, as the dataset is predominantly from the pre-vaccine era, the proportion of serotype 14 isolates was higher than 11A (four vs. one) within this genotype. The present dataset contained four GPSC10-24F strains, and have a predicted penicillin MIC of 0.5μg/mL.

### Comparison of core proteome sizes

The present study, therefore, attempted at constructing a pangenome for all the strains collected from India. Using an existing database with annotated genomes of 618 strains and relevant metadata, we created a pangenome using Panaroo. One of the main arguments of the tool is the family threshold which determines the clustering of related sequences based on their pairwise identity. While the default threshold of 0.7 is recommended, we tried to fine tune the settings by constructing a pangenome and observing the changes in core genome size when using different family thresholds. Using multiple family thresholds ranging from 0.7 to 1.0, we identified a threshold of 0.95 gives the largest core genome size while clustering proteins with high sequence homology. Thereafter, using this threshold value, the Indian Pneumococcal Pan-Genome (IPPG) is constructed with a total core genome size of 1615 genes, and 1534 of these were identified in >99% of the strains. The core genome size was slightly higher when compared with previous studies where the average core genome consisted of 500-1100 orthologous clusters [16].

However, we identified a similar pangenome size (4919 genes) as compared with ∼5000 genes in the pangenome. There were deviations identified from these core and pangenome sizes, like in a study from Thailand, the core genome was of 394 genes while the pangenome consisted of >13000 genes [18]. The above findings imply geographic diversity within the core and pangenome sizes. Using a Bayesian approach, the authors identified the core genome size of pneumococci can be around 851 genes [18].

Based on the existing GPS data, the most frequent GPSCs in India were GPSC1, GPSC2, GPSC8, GPSC9 and GPSC10. Pangenomes for these were constructed in a similar manner, and core genomes were extracted and re-annotated using Prokka. The core genome of GPSC10 is of least size, while GPSC1 had the highest size. We identified a decreasing trend in core genome size as the number of samples increased (Supplementary figure 2) – except for GPSC1 which had 52 genomes and the highest core genome size (Table 1).

### Antibiotic resistance genes

The GPSC1 core genome also harbored the tetracycline (tetM) and macrolide resistance (mefA-msrD) genes, while GPSC10 also had mefA-msrD genes. Presence of these antibiotic resistance genes indicates that these GPSC types can be frequently associated with resistance to tetracycline and macrolides. In fact, our study findings validate the widespread antibiotic resistance commonly associated with GPSC10 [4]. The reason for harboring these resistance genes and the impact on the bacterial metabolism can be explored further.

### Virulence genes

The proportion of virulence genes were higher among the GPSC1 core genome, and it lacked the capsular locus cps4A, cps4B, cps4C, cps4D, cps4F, cps4J, cps4K, and cps4L (Figure 4). Based on the GPS project findings, this GPSC1 is predominantly associated with serotype 19F and is most invasive [3]. In the present dataset, 44/52 GPSC1 isolates were of serotype 19F, among which 33 were from invasive specimens. However these genes were known to be conserved among the 19F serotype isolates [19]. One plausible explanation for this finding is the cutoff chosen for the core genome proportion. Including 95% of the strains, the gene has to be present among 49 isolates, and therefore may be missed due to this stringent cutoff. Other major virulence factors like pneumolysin (ply) responsible for host cell membrane pore formation, pneumococcal surface adhesin (psaA) for cell adhesion, metal ion binding protein pce (cbpE), fibronectin and RNA binding protein pavA, neuraminidase nanB, autolysin lytA, and hysA are detected among all the core genomes, indicating their indispensable role in pneumococcal pathogenesis. The hysA gene is specifically upregulated in the presence of hyaluronic acid (HA) and is involved in biofilm formation [20].

### Metabolic functions

Functional analysis using eggNOG database identified similar COG categories across all the core genomes (Figure 3), and most of the proteins were identified with category S (“unclassified”). Similar to our findings, Tonder et al identified category S as the most frequent category (21.7-24.1%) while major COG categories were J (11.9-15.7%), E (7.1-8.6%), and K (6.7-7.9%) [18]. In the present study, categories J (8.3-9.1%), P (6.5-8%) and F (6.8-7.9%) were predominant, and categories E and K were identified at 5.1-5.5% and 5.6-7.1% respectively (Supplementary File). Around 1.5-1.8% of the genes were of category V (defense mechanisms) and none of the core genomes had category X (mobilomes).

### Protein cluster and gene ontology analysis

These core proteomes were then compared with the IPPG core proteome to identify number of shared protein clusters using Orthovenn3. More than half of the clusters were shared among all the GPSC types and IPPG (figure 7), and GPSC1 and GPSC8 had no unique clusters within their core genomes. Around 21-24 clusters were uniquely identified among the remaining GPSCs and the IPPG. Around 462 clusters were shared among at least five proteomes, indicating an overall conserved proteome within the core genomes. When the GPSC core proteomes were analyzed separately, 1032 of the total 1693 clusters were shared and 405 clusters were not identified within the GPSC10 core genome (Supplementary Figure 3). The GPSCs 1, 2 and 9 shared a common proteome of 1619-1620 clusters, and were, therefore, highly similar in their core genome composition.

The Gene Ontology (GO) terms are assigned to more than 50% of the proteins within each core genome. Many genes had multiple GO terms associated with it, indicating complex functional roles carried out by these genes. Most (543/902) of the shared clusters are associated with metabolic functions (GO: 0008152), and a small proportion (41/902) of clusters are associated with hydrolase activity (GO: 0016787). Among the 405 clusters absent in GPSC10, the GO functional annotation terms predominated for ion binding (GO: 0043167, N=13), hydrolase (GO: 0016787, N=9), and transferase (GO: 0016740, N=7) activities.

**Supplementary figure 3:**
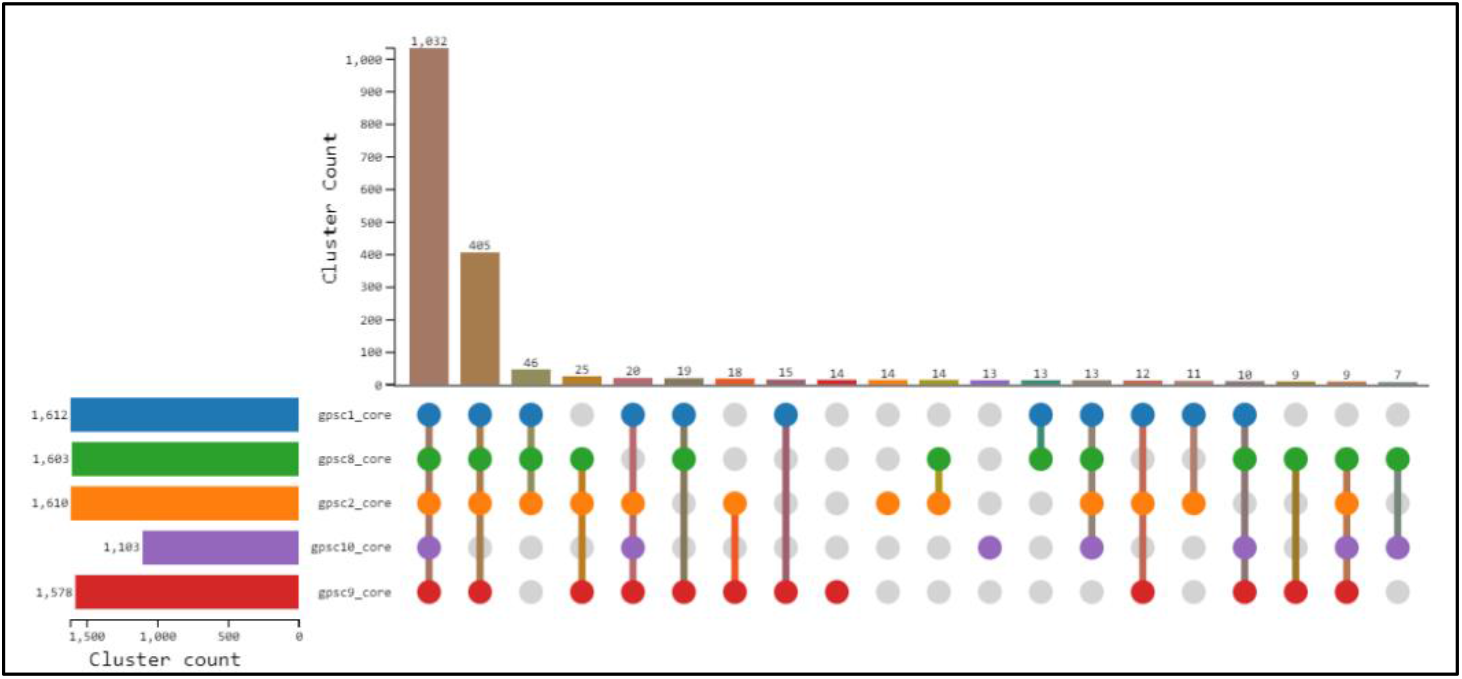
Core proteome clusters shared among the five GPSC types. 405 protein clusters were not identified in GPSC10 while it had 13 unique clusters.

In the present study, we therefore identified a large number of functionally conserved proteins and protein clusters among the major GPSC types. The presence of drug resistance genes within the core genomes of major GPSC that are also involved in invasive and non-vaccine serotypes raise further concerns regarding the long term evolutionary trajectories of these genotypes. Using core genome based phylogenetic analysis we identified a plausible serotype switching event from 14 to 11A within GPSC10 isolates.

Although the dataset is smaller in comparison with >1000 strains analyzed from other regions [16,18], our study is the first to identify and study the core genomic variations within the major GPSCs from India.

The functional roles have been predicted using COG and GO terms, and further analysis using protein interaction networks or metabolic networks is planned in the near future. The present study merits in understanding the genomic landscape of pneumococci by adopting the GPSC classifications. It is noteworthy that these classifications are further associated with metabolic genotypes, and can help in understanding strain replacement as observed in a recent study [5]. Using insights from these observations, one may identify the role of these common genes and genes unique to a GPSC type in bacterial metabolism and pathogenesis, and may aid in development of targeted therapies/vaccines. Some of the limitations of the present study are the low number (N=34) of isolates in the post-vaccine era, which prevented us from making any comparative analysis in genomic characteristics after the vaccine introduction.

## Conclusions

The Indian Pneumococcal Pan-Genome is of comparable size to that observed in other parts of the world, and is confirmed to be of”open” type. The presence of antibiotic resistance genes within the core genomes of GPSC1 and GPSC10 indicate the reason for the frequent antibiotic resistance among these genotypes. The most conserved virulence genes identified are involved in cell adhesion and biofilm formation, apart from other pathologic functions like autolysis and host membrane degradation. The GPSC10 shares the least number of genes within its core genome, indicating a widespread divergence of strains belonging to this genotype.

## Acknowledgements

The authors would like to acknowledge the efforts by the members of the Global Pneumococcal Sequencing project team in collecting and creating an open source database for analysis. The authors acknowledge Mr. Varun Shamanna (Bioinformatician, Central Research Lab, KIMS, Bengaluru) and Dr. K.L. Ravikumar (Chief, Central Research Lab, KIMS, Bengaluru) for providing training in genome analysis.

SRMP would like to acknowledge the Research Associate fellowship awarded by the Indian Council of Medical Research (ICMR) (file no: OMI-Fellowship/3/2020-ECD-I).

## Notes

https://data.monocle.sanger.ac.uk/

https://figshare.com/articles/dataset/GPSC_pangenome_analysis/24993954

### Competing Interest Statement

The authors have declared no competing interest.

### Summary of Updates

The figures were uploaded in line with the text to enhance clarity.

## References

1. Ministry of Health and Family Welfare, GoI. PCV Operational Guidelines [Internet]. Government of India; 2021. Available from: https://main.mohfw.gov.in/sites/default/files/PCV_Operational%20Guidelines_Jan%20%202021.pdf

2. GPS :: Global Pneumococcal Sequencing Project [Internet]. [cited 2022 Nov 14];Available from: https://www.pneumogen.net/gps/index.html

3. Gladstone RA, Lo SW, Lees JA, Croucher NJ, van Tonder AJ, Corander J, et al. International genomic definition of pneumococcal lineages, to contextualise disease, antibiotic resistance and vaccine impact. EBioMedicine 2019;43:338–46.

4. Lo SW, Mellor K, Cohen R, Alonso AR, Belman S, Kumar N, et al. Emergence of a multidrugresistant and virulent Streptococcus pneumoniae lineage mediates serotype replacement after PCV13: an international whole-genome sequencing study. Lancet Microbe 2022;3:e735–43.

5. Obolski U, Swarthout TD, Kalizang’oma A, Mwalukomo TS, Chan JM, Weight CM, et al. The metabolic, virulence and antimicrobial resistance profiles of colonising Streptococcus pneumoniae shift after PCV13 introduction in urban Malawi. Nat. Commun. 2023;14:7477.

6. Tonkin-Hill G, MacAlasdair N, Ruis C, Weimann A, Horesh G, Lees JA, et al. Producing polished prokaryotic pangenomes with the Panaroo pipeline. Genome Biol. 2020;21:180.

7. Minh BQ, Schmidt HA, Chernomor O, Schrempf D, Woodhams MD, von Haeseler A, et al. IQ-TREE 2: New Models and Efficient Methods for Phylogenetic Inference in the Genomic Era. Mol. Biol. Evol. 2020;37:1530–4.

8. Page AJ, Taylor B, Delaney AJ, Soares J, Seemann T, Keane JA, et al. SNP-sites: rapid efficient extraction of SNPs from multi-FASTA alignments. Microb. Genomics 2016;2:e000056.

9. Seemann T. Prokka: rapid prokaryotic genome annotation. Bioinforma. Oxf. Engl. 2014;30:2068–9.

10. Sun J, Lu F, Luo Y, Bie L, Xu L, Wang Y. OrthoVenn3: an integrated platform for exploring and visualizing orthologous data across genomes. Nucleic Acids Res. 2023;51:W397–403.

11. Huerta-Cepas J, Szklarczyk D, Heller D, Hernández-Plaza A, Forslund SK, Cook H, et al. eggNOG 5.0: a hierarchical, functionally and phylogenetically annotated orthology resource based on 5090 organisms and 2502 viruses. Nucleic Acids Res. 2019;47:D309–14.

12. Seemann T. tseemann/abricate [Internet]. 2024 [cited 2024 Jan 12];Available from: https://github.com/tseemann/abricate

13. Feldgarden M, Brover V, Haft DH, Prasad AB, Slotta DJ, Tolstoy I, et al. Validating the AMRFinder Tool and Resistance Gene Database by Using Antimicrobial Resistance Genotype-Phenotype Correlations in a Collection of Isolates. Antimicrob. Agents Chemother. 2019;63:e00483–19.

14. Chen L, Zheng D, Liu B, Yang J, Jin Q. VFDB 2016: hierarchical and refined dataset for big data analysis—10 years on. Nucleic Acids Res. 2016;44:D694–7.

15. Shannon P, Markiel A, Ozier O, Baliga NS, Wang JT, Ramage D, et al. Cytoscape: a software environment for integrated models of biomolecular interaction networks. Genome Res. 2003;13:2498–504.

16. Hiller NL, Sá-Leão R. Puzzling Over the Pneumococcal Pangenome. Front. Microbiol. 2018;9:2580

17. Nagaraj G, Govindan V, Ganaie F, Venkatesha VT, Hawkins PA, Gladstone RA, et al. Streptococcus pneumoniae genomic datasets from an Indian population describing pre-vaccine evolutionary epidemiology using a whole genome sequencing approach. Microb. Genomics 2021;7:000645.

18. Andries J van Tonder, James E Bray, Keith A Jolley, Sigríður J Quirk, Gunnsteinn Haraldsson, Martin CJ Maiden, et al. Heterogeneity among estimates of the core genome and pan-genome in different pneumococcal populations. bioRxiv 2017;133991.

19. Morona JK, Morona R, Paton JC. Analysis of the 5′ Portion of the Type 19A Capsule Locus Identifies Two Classes of cpsC, cpsD, andcpsE Genes in Streptococcus pneumoniae. J. Bacteriol. 1999;181:3599–605.

20. Yadav MK, Chae SW, Park K, Song JJ. Hyaluronic acid derived from other streptococci supports Streptococcus pneumoniae in vitro biofilm formation. BioMed Res. Int. 2013;2013:690217.

